# Mutation-centric Network Construction using Long-Range Interactions

**DOI:** 10.64898/2026.03.16.712158

**Authors:** Ramal Hüseynov, Burçak Otlu

## Abstract

Somatic mutations can alter normal cells and lead to cancer development. Yet distinguishing functional driver mutations from neutral passenger mutations remains a significant challenge. Traditional genomic tools often prioritize linear overlap searches, failing to capture the complex, three-dimensional regulatory environment of the genome. We present a graph-based framework, MutationNetwork, for constructing mutation-centric networks by integrating long-range intrachromosomal interactions with local genomic overlaps. Our method utilizes a unique positive and negative indexing scheme to represent interacting genomic intervals as nodes. By encoding both interactions and overlaps as edges, we enable constant-time retrieval of complex relationship data. By iteratively expanding the graph from a seed mutation, we can quantify a mutation’s influence on the genomic landscape and assess its proximity to genes. We applied this framework to a dataset of 560 breast cancer whole-genome sequences, focusing on Triple-Negative Breast Cancer (TNBC) and Luminal A subtypes. Our results demonstrate that the generated mutation embeddings successfully cluster samples according to their biological subtypes, with the highest classification performance achieved at specific ranges. This approach provides a comprehensive view of mutation impact, offering a scalable solution for cancer patient stratification and the prioritization of potential non-coding driver mutations by assessing their network-level impact.

**Availability and implementation:** The source code is available at https://github.com/Ramalh/MutationNetwork

## Introduction

Long-range genomic interactions extend beyond simple spatial packing, forming complex regulatory networks where distant regions modulate one another’s function (Dekker 2003]). While many somatic mutations occur within distal regulatory elements, identifying their functional targets is challenging because these elements typically regulate genes through intricate 3D chromatin architecture rather than simple linear proximity. Large-scale initiatives like ENCODE have systematically mapped these associations, producing ‘loop files’ that catalog interacting genomic regions by their precise coordinates (Luo et al. [2020]). However, existing genomic interval tools, including BEDTools (Quinlan and Hall [2010]), PyRanges (Stovner and Sætrom [2019]), and various IntervalTree implementations, are not designed to model the multi-step relationships linking mutations to distal genes through such chromatin interactions. Because these loop files integrate both positional and chromosomal interaction data, they are critical for determining whether a mutation’s impact propagates to distal genomic intervals.

To capture these dependencies, the genome can be modeled as a complex network where interacting genomic intervals are represented as nodes, and physical overlaps or functional interactions serve as edges. Although existing tools are highly optimized for 1D interval arithmetic and overlap detection, they lack the native architecture required to integrate and traverse heterogeneous relationships within a unified graph structure, which is essential for assessing the full regulatory impact of a mutation. To overcome these limitations, we introduce a method that maps all long-range interacting intervals onto an array-based structure while preserving critical interaction and overlap data. Using arrays and sets to manage these relationships, our approach eliminates the overhead of recurring interval querying, significantly accelerating the assessment of mutation impacts.

## Methods

Input data are provided in the BEDPE format, where each row defines a discrete pairwise interaction between two genomic intervals. While each interval is represented by a single pair of coordinates, an interval involved in multiple interactions appears across multiple rows. Our framework identifies these multi-contact points by detecting coordinate overlaps, thereby integrating discrete genomic intervals into a unified, mutation-centric network (Figure 1A).

**Fig. 1.**
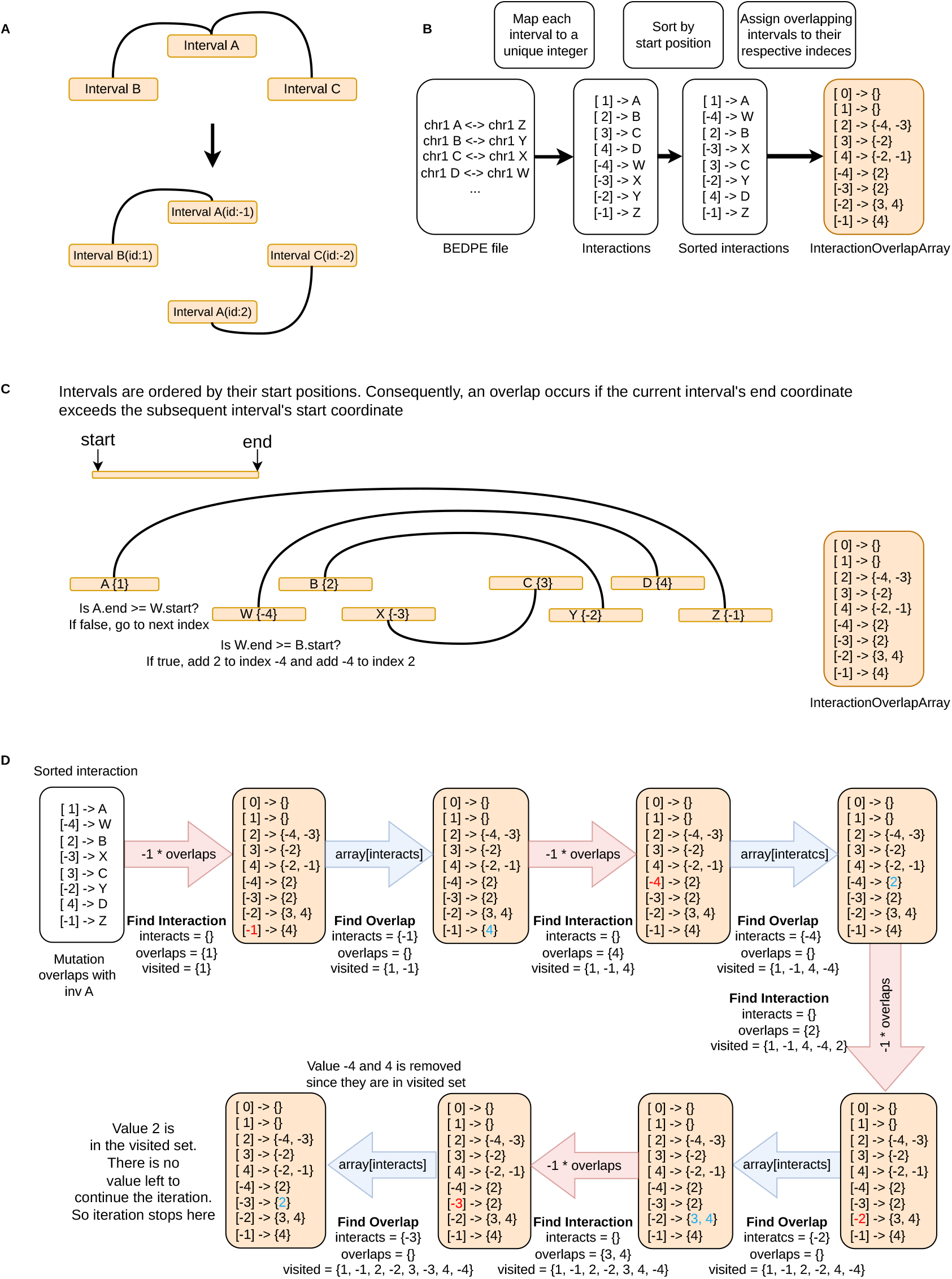
**(A)** Multi-interacting regions represented as distinct intervals on separate lines. **(B)** Construction of the `InteractionOverlapArray` from a BEDPE file. **(C)** Overlap detection using sorted intervals. **(D)** Graph construction utilizing the `InteractionOverlapArray` to retrieve identified overlaps and interactions.

The proposed method consists of three primary steps: (i) transforming the BEDPE file into an array of genomic intervals (Figure 1B), (ii) identifying all overlapping intervals and mapping them to their corresponding array indices (Figure 1C), and (iii) performing a graph traversal starting from the mutation site, iteratively incorporating all connected intervals to define the final network (Figure 1D).

The first step involves preprocessing the BEDPE file to facilitate rapid interaction retrieval. Each of the *L* pairwise interactions is assigned a unique positive integer *i* ∈ {1, …, *L*}, where *L* represents the total number of rows in the BEDPE file. We then deconstruct the BEDPE structure into a one-dimensional array of 2*L* + 1 entries, where the ‘left’ interval of interaction *i* is stored at index *i*, and the ‘right’ interval is stored at index *−i* (Figure 1B). Index 0 is reserved as a dummy entry. This symmetric indexing scheme enables direct constant-time access to any interaction partner; for instance, the partner of the interval at index 5 is immediately retrieved at index *−*5, and vice versa.

### InteractionOverlapArray Construction

The second step captures overlap relationships by initializing an array of empty sets, denoted as the `InteractionOverlapArray`, which corresponds to the 2*L* + 1 indices defined in the previous step. Following a start coordinate-based sort of the intervals, each entry is compared with its neighbors to identify overlaps. When an overlap is detected, for example, between the interval at index *−*4 and the interval at index 2, the corresponding indices are cross-referenced: *−*4 is added to the set at index 2, and 2 is added to the set at index *−*4 (Figure 1C). Because the genomic intervals and their pairwise interactions remain static for a given BEDPE file, this array provides a persistent, constant-time lookup for all topological relationships.

The construction of the `InteractionOverlapArray` is detailed in Algorithm 1. The process begins with the linearization of the BEDPE file into a one-dimensional array, `InvArray` (Lines 1–4). In Line 1, the coordinates of the left set of genomic intervals are mapped to the first *L* entries. Line 2 assigns these entries unique positive identifiers from 1 to *L*. To maintain a symmetrical mapping, Line 3 populates the remaining *L* entries with the right set of intervals in reverse order. Line 4 then assigns these intervals corresponding negative identifiers. This indexing scheme ensures that both left and right intervals of the same interaction share a common absolute ID, allowing the algorithm to track the interacting genomic intervals throughout the computation. Lines 5–6 initialize the `InteractionOverlapArray` as an adjacency list of size 2*L* + 1, where each entry is an empty set. To facilitate an efficient search, Line 7 sorts `InvArray` by start position in ascending order. While BEDPE files are frequently pre-sorted, this explicit sort guarantees the correctness of the subsequent sweep-line scan. The scanning phase (Lines 8–18) leverages the sorted structure to identify physical overlaps in genomic space. For each interval *i*, the algorithm evaluates succeeding intervals *j*. The condition in Line 13, checking if the end of interval *i* is greater than or equal to the start of interval *j*, determines a physical overlap. When an overlap is detected, the formerly stored identifiers are added to the respective sets in `InteractionOverlapArray` (Lines 14–15). The use of a set data structure inherently handles redundancy; overlap relationships are recorded only once. Algorithm 1 runs in *O*(*L* log *L* + *K*) time, where *L* is the number of interactions and *K* is the number of overlapping pairs. The sort dominates at *O*(*L* log *L*), and the sweep line loop adds *O*(*K*) for processing overlaps. The algorithm is output-sensitive, fast when overlaps are sparse (typical in genomic data), and worst-case *O*(*L*^2^) when all intervals overlap.

### Mutation-Centric Graph Construction

The construction of the mutation-centric network is detailed in Algorithm 2. This procedure builds a localized graph G by treating the `InteractionOverlapArray` as an adjacency list and performing a Breadth-First Search (BFS) to traverse and construct a subgraph of overlaps and interactions. The algorithm initializes a graph G using the NetworkX library (Hagberg et al. [2008]), with the mutation site serving as the central “seed” node. Lines 2–6 perform an initial check to determine if the mutation has any overlaps. If no overlaps are found, the algorithm terminates and returns an empty graph. For mutations with valid overlaps, Lines 7–10 establish the first layer of connectivity. This stage defines the interacting partners (represented by the negation *−i*). This mathematical negation is a coding convention used to distinguish between a physical genomic segment ID and its functional interacting partner. The core of the algorithm (Lines 11–22) expands the graph through a standard BFS “frontier” expansion. The network grows outward through cumulative layers of connectivity, where each range *k* defines the depth of the included relationships. For instance, a range 0 restricts the subgraph to the mutation node itself, whereas a range 1 includes the mutation node and its immediate overlapping genomic intervals. Range 2 extends to include the functional interaction partners of those overlapping intervals. For subsequent ranges (*k ≥* 3), the process repeats this alternating pattern, successively incorporating further layers of spatial overlaps and functional interactions. As the algorithm “walks” through these connected components, it populates G with both spatial and functional edges. To prevent infinite loops or redundant processing, a visited set (Line 21) tracks processed nodes, ensuring the algorithm remains efficient. Because the algorithm only visits each connected node and edge once, it achieves a time complexity of *O*(*V* + *E*), where *V* is the number of nodes and *E* is the number of edges. This localized approach is significantly more efficient than a global genome scan. Biologically, this structure captures how a mutation may influence nearby genomic regions through overlaps, which can subsequently connect to distal genomic regions via chromatin interactions. This traversal captures higher-order genomic relationships within a user-defined distance, yielding a connected graph where nodes are annotated with relevant biological attributes, such as overlapping gene features.

### Feature Matrix Generation

Following the graph construction, these mutation-centric subgraphs are vectorized to facilitate downstream analysis. For each mutation, a binary gene-presence vector of dimension *N* is generated, where *N* represents the total number of genes in the analysis. For a specified range k, the *i*^*th*^ element of the binary vector is set to 1 if gene *g*_*i*_ is contained within the set of nodes belonging to the mutation-centric subgraph (Figure 2). Because a single sample typically harbors multiple mutations, the individual vectors for that sample are consolidated into a single representative profile using an element-wise maximum operation. This process effectively identifies all genes “reachable” from any mutation in the sample at the specified depth. Finally, these consolidated vectors are aggregated into an *M × N* feature matrix for each specified range *k*, where *M* is the total number of samples, providing a structured representation of the functional and spatial impact of the mutational landscape across the cohort.

**Fig. 2.**
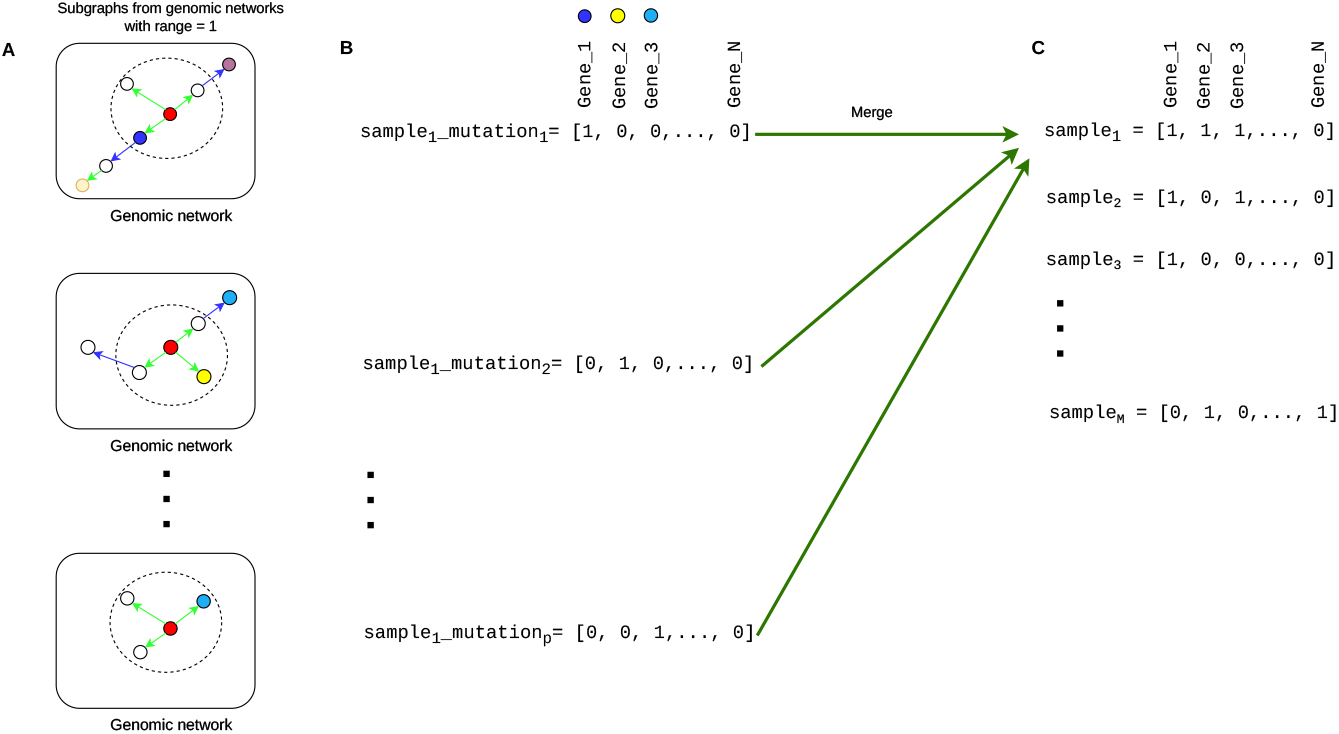
Overview of Mutation-centric Network. **(A)** For each mutation within a sample, a mutation-centric subgraph is constructed based on a defined range *k*. Only nodes within a shortest-path distance *d ≤ k* from the seed mutation (the central node) are retained, while distal nodes are pruned. **(B)** For a given mutation, a binary row vector is generated where the *i*^*th*^ element is set to 1 if the corresponding gene *g*_*i*_ is present within the subgraph. **(C)** To represent the entire sample, individual mutation vectors are aggregated into a single representative row vector using an element-wise *maximum* (OR) operation. This results in a final binary feature matrix of dimension *M × N*, where *M* is the total number of samples and *N* is the number of genes in the analysis.

#### Algorithm 1

Creating `InteractionOverlapArray` from a BEDPE file with time complexity 𝒪(*L* log *L* + *K*), where *L* is the number of interactions and *K* is the number of overlapping pairs.

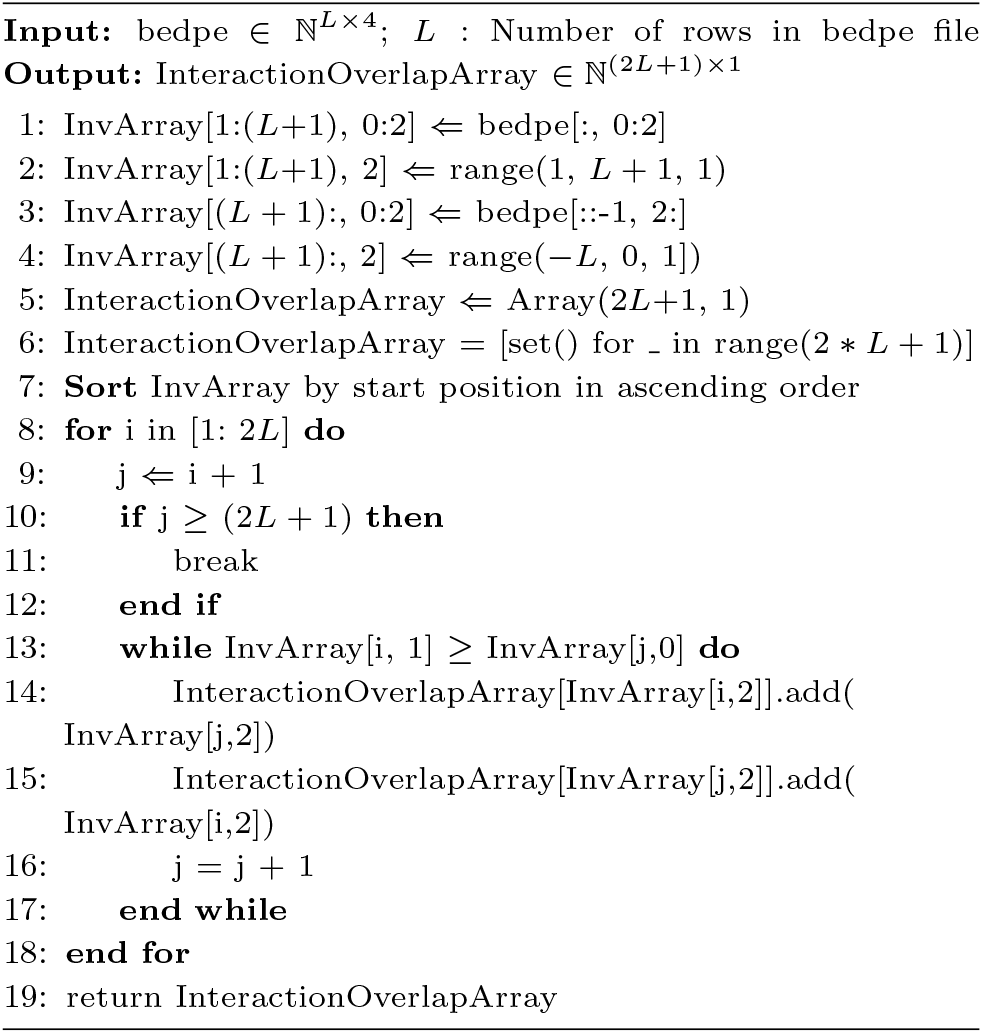

#### Algorithm 2

Create graph G with interactions and overlaps for the given mutation, array and range *r* with time complexity *O*(*V* + *E*) where *V* is the number of nodes and *E* is the number of edges.

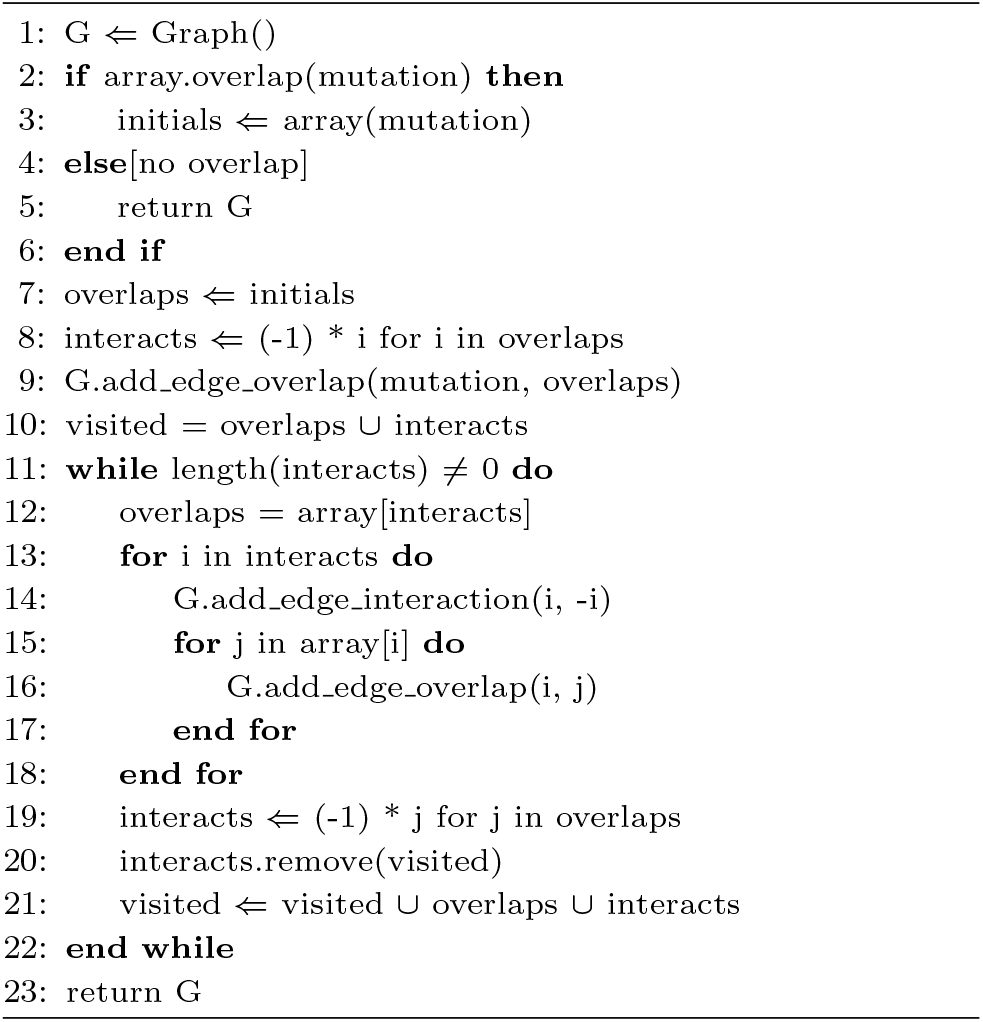

### Dataset

The experimental dataset was aggregated from multiple genomic sources. We obtained somatic mutation data from 560 breast cancer whole-genome sequences reported by Nik-Zainal et al. [2016]. The original study provides clinicopathological annotations, including estrogen receptor (ER), progesterone receptor (PR), and human epidermal growth factor receptor 2 (HER2) status. Based on these profiles, we categorized the samples into two distinct molecular subtypes for downstream analysis: Luminal A (ER+, PR+, HER2-) and Triple-Negative Breast Cancer (TNBC; ER-, PR-, HER2-) (Orrantia-Borunda et al. [2022]). We focused on these two subtypes as they represent biologically divergent entities and comprise the majority of the available cohort. The final cohort comprised 276 Luminal A and 163 TNBC samples, totaling 439 cases. Chromatin interaction data (BEDPE format) were sourced from the ENCODE project (Experiment ID: ENCSR059HDE), specifically utilizing loops derived from breast cancer cell lines to ensure tissue specificity. Finally, gene annotations were obtained from the GENCODE v47 basic gene set, encompassing a total of 77,114 genes. To ensure data integrity, entries with duplicate gene names were excluded.

## Results

To evaluate performance, we implemented an alternative workflow using PyRanges as a baseline for comparative analysis. We quantitatively assessed the execution time of our array-based approach compared to PyRanges for detecting interval overlaps. As illustrated in Figure 1, the two workflows share common steps; however, they diverge in how they manage interval interactions. While the MutationNetwork framework utilizes the array-based structures described in Figures 1B and 1C to catalog all interaction and overlap data, the alternative approach replaces these specific steps with PyRanges-based queries. Both methods then proceed to the final step (Figure 1D) using the same `InteractionOverlapArray`. The results of this temporal comparison are summarized in Table 1 and Figure 3.

**Table 1.** Performance Benchmarking: Execution Times for MutationNetwork vs. PyRanges using BEDPE Dataset ENCFF597SQA (Seconds).

| File | MutationNetwork | MutationNetwork+Pyranges |
| --- | --- | --- |
| PD9844a | 259 | 925.5 |
| PD18101a | 484.2 | 1149.0 |
| PD11767a | 582.0 | 1261.0 |
| PD22251a | 880.9 | 1548.8 |
| PD9541a | 1479.9 | 2153.4 |
| PD24199a | 1550.3 | 2226.3 |
| PD6406a | 2935.3 | 3632.3 |
| PD24202a | 4428.8 | 5126.0 |
| PD3945a | 5894.3 | 6532.8 |
| PD9604a | 6449.1 | 7112.5 |
| PD8832a | 13994.1 | 14762.6 |

**Fig. 3.**
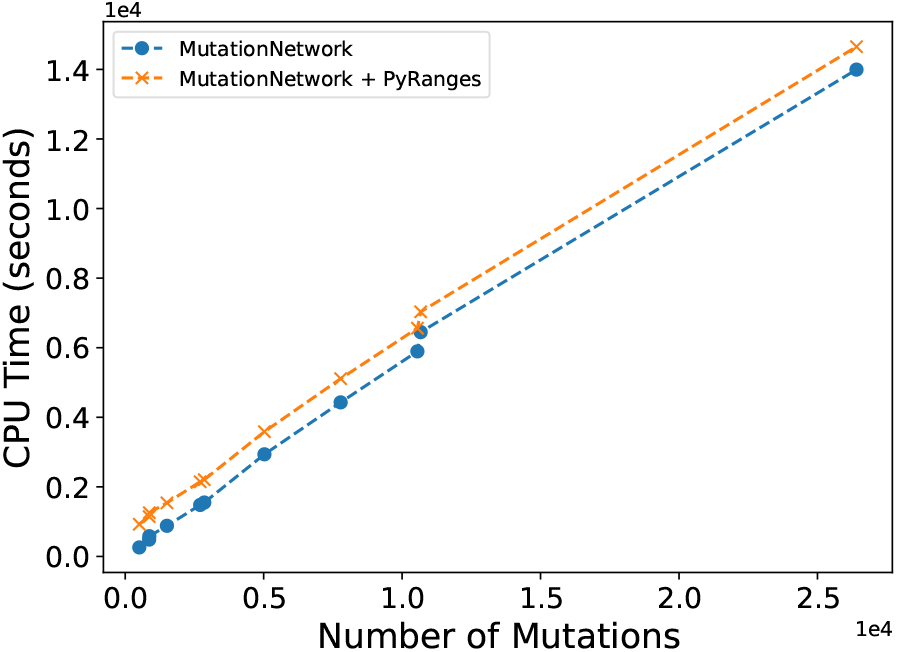
Execution time comparison between MutationNetwork and PyRanges.

For a given set of samples, a chromatin loop file, and a specified range, the pipeline generates a binary feature matrix where rows represent samples and columns represent gene features. To investigate whether samples cluster according to their affected genes and if these clusters correlate with known breast cancer molecular subtypes, we implemented a dimensionality reduction and clustering workflow. Given the initial high-dimensional space (*∼* 80,000 genes), we first reduced the feature space down to 250 components using Truncated Singular Value Decomposition (SVD) (Hansen [1987]). Truncated SVD was implemented by scikit-learn library (Pedregosa et al. [2011]). This was followed by Uniform Manifold Approximation and Projection (UMAP) to project the data into 15 components, capturing non-linear relationships within the dataset (McInnes et al. [2018]). The quality of these reductions was monitored via explained variance for SVD and trustworthiness for UMAP, maintaining thresholds above 0.75 and 0.60, respectively, across all ranges.

To evaluate the biological signal captured at different ranges, we performed hierarchical clustering on the reduced feature matrices using cosine similarity. By partitioning the resulting dendrograms at the primary branch level, we assigned samples to predicted ‘TNBC’ or ‘Luminal A’ classes and validated these against known molecular subtype annotations. As shown in Figure 4, the dendrograms for ranges 4, 5, and 14, contrasted against a range 0 baseline representing only direct mutation sites (Figure 4A), reveal two highly distinct clusters. The significant inter-cluster separation and high intra-cluster cohesion at these ranges suggest that chromatin-loop-derived features provide a robust biological signal for distinguishing molecular subtypes (Figure 4B-D).

**Fig. 4.**
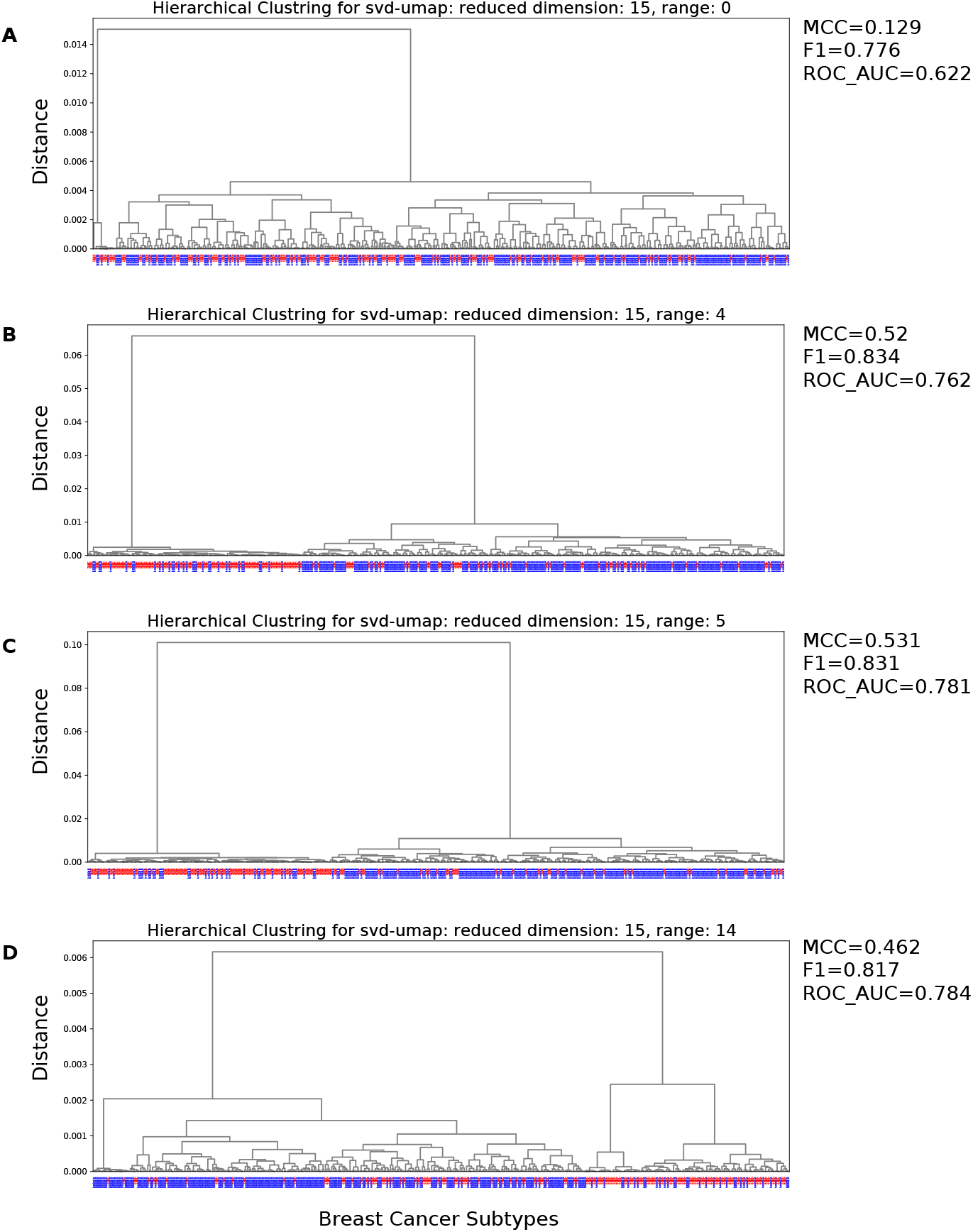
Hierarchical clustering of breast cancer samples across topological ranges. **(A** Baseline dendrogram at range *k* = 0 (direct mutation-to-gene overlaps), demonstrating minimal subtype separation. **(B-D)** Dendrograms for ranges *k* = 4, *k* = 5, and *k* = 14 where the integration of long-range chromatin interactions reveals a distinct bifurcation into predicted ‘TNBC’ (red) and ‘Luminal A’ (blue) molecular subtypes. Clustering was performed using cosine similarity on reduced feature matrices. The high intra-cluster cohesion and marked inter-cluster divergence at these depths indicate that the mutation-centric network effectively captures the three-dimensional regulatory landscape necessary for high-resolution subtype discrimination.

To assess classification performance, we calculated metrics including the Matthews Correlation Coefficient (MCC), F1-score, and Area Under the ROC Curve (AUC-ROC) (Figure 5). The ROC curves were generated using a distance-to-centroid scoring method. Performance metrics (MCC, F1-score, and AUC-ROC) across these clusters demonstrate a clear peak in classification performance at ranges 4 and 5, with a distinct secondary peak observed between ranges 13 and 15. (Figure 5). Beyond these ranges, performance metrics converge, suggesting that the predictive signal becomes saturated.

**Fig. 5.**
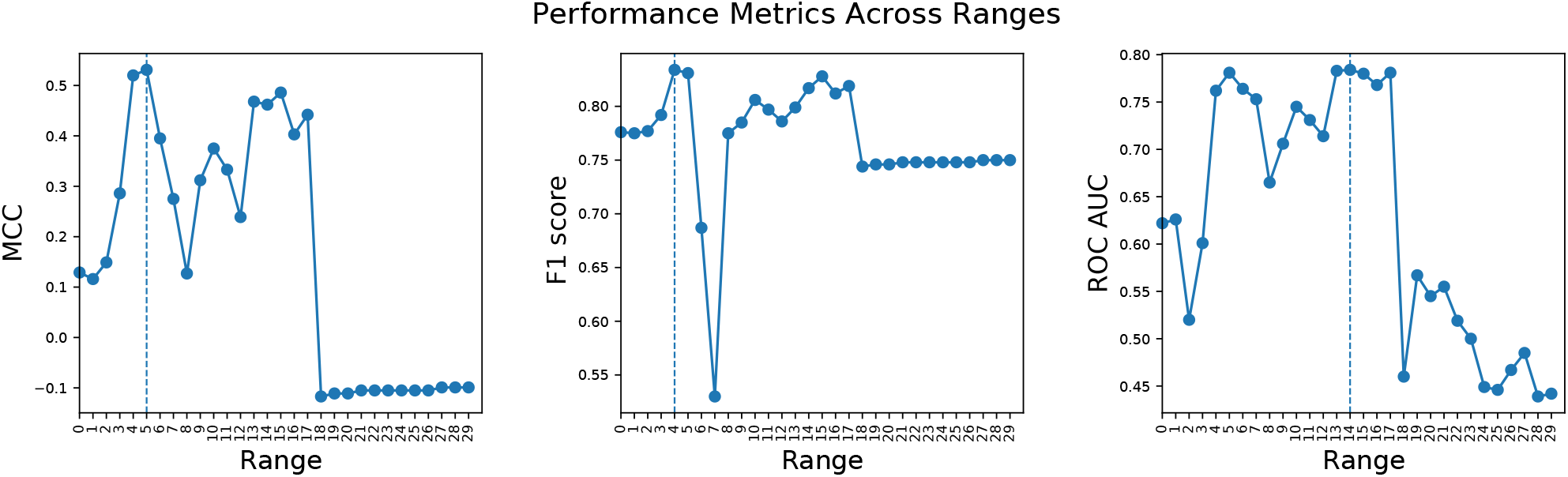
Classification performance across different ranges. The plot displays the Matthews Correlation Coefficient (MCC), F1-score, and Area Under the ROC Curve (AUC-ROC). A clear peak in classification performance is observed at range 4 and range 5, followed by a convergence of metrics at higher ranges, suggesting a saturation of the predictive signal. ROC curves were generated using a distance-to-centroid scoring method.

## Conclusion

This study introduces a novel graph-based framework for constructing mutation-centric networks that integrate 1D genomic overlaps with 3D long-range chromosomal interactions. By utilizing a symmetric indexing scheme and an array-based architecture, our approach achieves constant-time retrieval of complex genomic relationships, significantly outperforming traditional interval-based tools in network traversal tasks. Applying this framework to 560 breast cancer whole-genome sequences, we demonstrated that the functional impact of mutations extends beyond linear proximity. Mutation embeddings effectively cluster samples by biological subtype, with peak classification performance achieved at ranges 4 and 5, followed by a localized increase around range 14. This suggests that these specific network depths capture the most relevant regulatory disruptions in Luminal A and TNBC. Ultimately, this scalable method provides a robust pipeline for prioritizing non-coding driver mutations and enhancing patient stratification through a systems-level view of the cancer genome.

## Competing interests

No competing interest is declared.

## Author contributions statement

R.H. developed the Python code and wrote the draft of the manuscript. B.O. conceived the idea, supervised the overall development of the code, and wrote the manuscript. All authors read and approved the final manuscript.

## Acknowledgments

This work was supported by the Akademik Gelişim Programı (AGEP) under Grant AGEP-704-2024-11495 by the Middle East Technical University. The numerical calculations reported in this paper were fully performed at TUBITAK ULAKBIM, High Performance and Grid Computing Center (TRUBA resources). The authors also thank Sana Basharat and Ahmet Emre Özdemir for testing the tool and for helpful discussions, as well as Hüseyin Hilmi Kilinç for reading the draft of the manuscript and for providing feedback.

## Notes

### Competing Interest Statement

The authors have declared no competing interest.

